# Newborn chicks show inherited variability in early social predispositions for hen-like stimuli

**DOI:** 10.1101/071456

**Authors:** Elisabetta Versace, Ilaria Fracasso, Gabriele Baldan, Antonella Dalle Zotte, Giorgio Vallortigara

## Abstract

Predispositions of newborn vertebrates to preferentially attend to living beings and learn about them are pervasive. Their disturbance (e.g. in human neonates at risk for autism), may compromise the proper development of a social brain. The genetic bases of such predispositions are unknown. Here we take advantage of well-known visual preferences exhibited by newly-hatched domestic chicks (*Gallus gallus*) for the head/neck region of their mother hen, to investigate the presence of segregating variation in the predispositions to approach a stuffed hen *vs.* a scrambled version of it. We compared the spontaneous preferences of three different breeds that have been maintained genetically isolated for at least eighteen years and identically raised in the same farm. Visually-naïve chicks of all the three tested breeds (Padovana, Polverara and Robusta maculata) showed the same initial preference for the predisposed stimulus, suggesting that the direction of the initial preference might be genetically fixed. A few minutes later though, striking differences emerged between breeds, which could indicate early different strategies of dealing with affiliative objects: while the Polverara breed maintained a constant preference across the entire test, the Padovana and Robusta breeds progressively explored the alternative stimulus more. We argue that exploration of novelty might help chicks to look for responsive parental objects and to form a more structured representation of the mother hen. We hence documented the presence of inherited genetic variability for early social predisposition in interaction with environmental stimuli.

## Introduction

Attending to animate stimuli since the beginning of life can be adaptive for species that require early social care. In social species, mechanisms that help individuals orienting towards animate objects soon after birth have been identified in young chicks, human and non-human primates [reviewed in 1]. Spontaneous preferences for cues associated with potential social partners include biases for attending to face-like configurations [2–4], biological *vs.* rigid motion [5–7], changes of speed [8] and self-propelled objects [9,10]. Recently it has been shown that neonates at high familiar risk of developing Autism Spectrum Disorders exhibit significantly weaker preferences for attending biological motion and face-like stimuli compared to control neonates [11]. Some of the stimuli used for testing human neonates have been first investigated in non-human models [5,12], showing the relevance and translational value of studies on early predispositions for animate objects in biomedical research. It is not known, though, to which extent early predispositions have a genetic basis. The chick of the domestic fowl (*Gallus gallus*) is a convenient subject to address this issue, due to the well-known presence of predispositions for orienting towards animate objects [1,13,14], the ease of control-rear chicks until the testing time, and the presence of breeds that have been maintained genetically separated during domestication [15,16]. Observing differences in early predispositions between chicken breeds would indicate the presence of natural genetic variability for this trait. In this study we investigated the spontaneous preferences of visually naïve chicks of different breeds for approaching a stuffed hen *vs.* a scrambled-hen (a stuffed hen whose parts were attached on the sides of a box in a scrambled order, see Figure 1). Spontaneous preferences for a stuffed hen have been repeatedly documented in broiler chicks: [12,17,18], and depending on the integrity of the neck and face region [3], one of the target of predisposed behaviours in human neonates [19–21]. In the chicks’ literature [12,18], the average preferences for the hen stimulus varied between 59 and 73%, but the average results include chicks with a strong preference for the stuffed hen as well as chicks that preferred the scrambled-hen. The source of the observed individual variability is unknown. To investigate the role of the genetic components in determining early preferences for hen-like stimuli, we compared the spontaneous preferences of three genetically isolated breeds identically raised in the same farm. These breeds belong to a conservation project (Co.Va [22]) and have been maintained genetically separated for more than eighteen years, so that there is low level of admixture between them [16]: Padovana (isolated since 1987), Polverara (isolated since 1998) and Robusta maculata (isolated since 1998). The genetic differentiation and phylogenetic distance between these breeds had been previously documented [16,23–26] but had never been linked to predispositions for affiliative responses.

**Figure 1.**
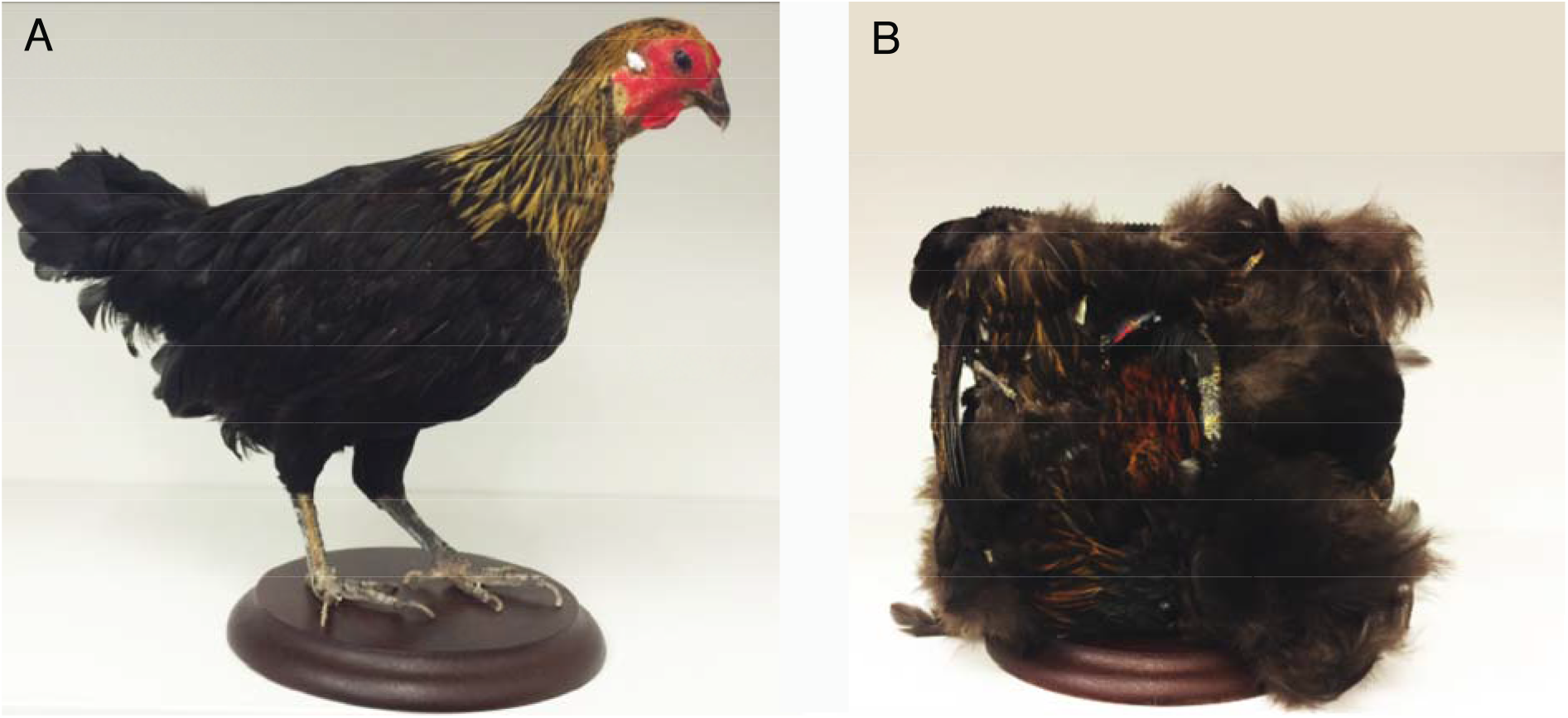
Pictures of the stimuli used: (A) Stuffed hen. (B) Scrambled hen (a stuffed hen whose parts were attached on the sides of a box in a scrambled order).

## Materials and methods

### Ethics statement

All experiments comply with the current Italian and European Community laws for the ethical treatment of animals and the experimental procedures were approved by the Ethical Committee of University of Trento and licensed by the Italian Health Ministry (permit number 1138/2015 PR).

### Breeds and conservation scheme

We investigated three chicken breeds that entered the Co.Va conservation project [22] since 2006: Padovana, Polverara and Robusta maculata.

Historical records suggest that the Padovana breed has been introduced in Italy from Poland more than seven centuries ago [22]. Until the beginning of the XX century, Padovana and Polverara breeds were confused, more than likely because both breeds are similar and have a tuft of feathers on their head (although in the case of Padovana it is more pronounced due to a skull ernia). The local market’s main interest is the meat production from Padovana and Polverara breeds. Strains of the Padovana breed include black, white, gold, silver, and buff coloured plumage, whereas the Polverara’s include black and white plumage [24]. In our study we considered individuals from gold, silver and buff Padovana, white and black Polverara, since previous studies revealed high homogeneity within these breeds [16]. The Robusta maculata breed was developed in 1965 at the Rovigo Experiment Station from crosses between Tawny Orpingtons and White Americans [22]. This breed was selected to provide both eggs and meat.

Zanetti et al. [16] documented genetic isolation (low level of admixture) between the investigated breeds, and a closer phylogenetic relationship between Padovana and Polverara, which are also more similar at phenotypic level compared to Robusta. Similarly, De Marchi et al. [24] have observed a close genetic relationship between Padovana and Polverara.

For all flocks, the breeding and conservation scheme aimed at increasing the number of pure breed animals with no gene flow between breeds, and maintaining genetic variability within the breed. In 2010, the population size has been estimated as ∼2000 for Padovana and ∼1500 for both Robusta maculata and Polverara [16]. The reproduction season starts at the end of January and birds hatch from February to June. New male and female reproducers representative of the breed are selected in October. Our experiment was conducted in 2016: we selected 40-45 females and 15 males for each variety of the Padovana breed buff and gold plumage; 20 females and 7-8 males for each variety of the Padovana breed of silver, black and white plumage; 23 females and 2 males for the Robusta breed; 20 females and 7 males for each variety of the Polverara breed white and black plumage. In January, males of each breed were divided in pairs and rotated every month among groups of 20-22 females. For all breeds, the reproducers were kept in enclosures with an indoor (3×4.5 m) and an outdoor part (3×15 m), with 2 males and 20-22 females each, fed with poultry feed Progeo (Reggio Emilia, Italy) *ad libitum*, in a light:dark regime of 15:9 hours.

### Subjects

Overall we tested 91 naïve domestic chicks (*Gallus gallus*): 31 Padovana, 31 Polverara and 29 Robusta maculata individuals (Figure 2). One Padovana chick did not move during the test and was excluded from further analyses. Eggs were obtained in 7 batches from the Agricultural High School “Duca degli Abruzzi” (Padova, Italy), which is pursuing the Co.Va conservation program for the maintenance of local biodiversity [22] described above.

**Figure 2.**
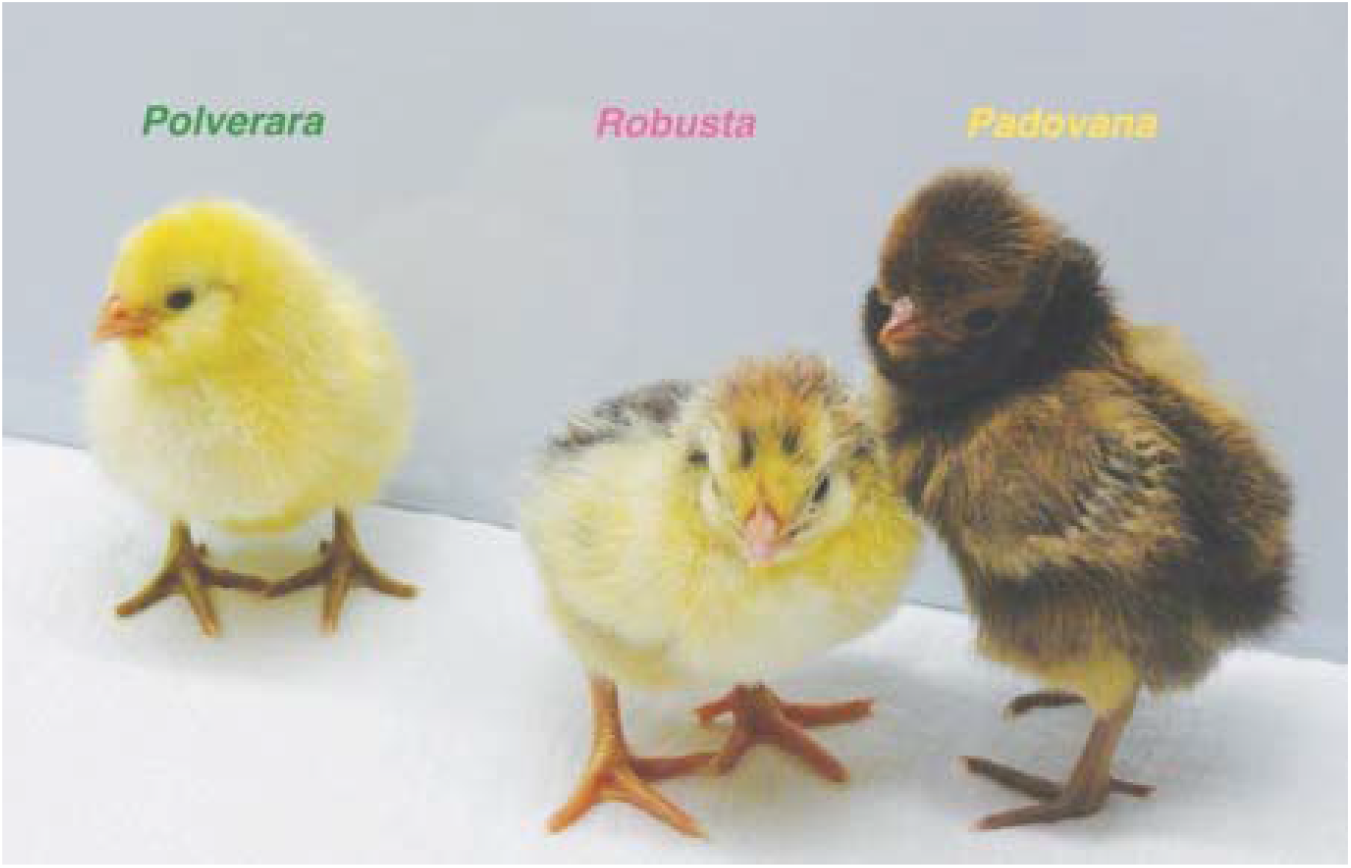
Chicks of the three investigated chicken breeds after the test, from the left: Polverara, Robusta and Padovana.

Eggs were incubated in darkness at 37.7 °C at 40% humidity for 17 days, then humidity was increased to 60% during the last three days of incubation. Twenty-four hours after hatching, chicks were transferred in individual compartments to an incubator at 33 °C (8×10×14 cm) and exposed to an unspecific acoustic stimulation (this procedure enhances predisposed preferences and had been performed in similar experiments by [18,27]). We used the same aspecific acoustic stimulation presented by [18], which consists in intermittent non-repeating rhythmic music segments played by a loudspeaker for 180 minutes overall. After the acoustic stimulation, chicks were maintained in individual compartments within a dark incubator until test. Test occurred 24 (+/- 8 hours) after acoustic stimulation, when chicks were 40 to 56 hour-old. Chicks were constantly kept in darkness until the moment of test.

### Test apparatus

The enclosure used for the test was 150 cm long, 46 cm wide, 45 cm high, with a running wheel (32 cm diameter, 13 cm large, covered with 1 cm of opaque foam on both sides) located in the middle of the apparatus. As stimuli we presented a stuffed hen and a scrambled-hen, which was prepared by scrambling the pieces of a stuffed hen disrupting the configuration of the head and neck (see Figure 1). Stimuli were located at the opposite sides of the apparatus, on two rotating platforms (20 rotations/minute). The stuffed hen closely resembled the jungle fowl hen used in previous studies [3,18]. The position of the stuffed hen and scrambled hen in the apparatus was counterbalanced between subjects. The stimuli were illuminated by an above light (40 W warm light) that diffused through a semi-transparent white plastic sheet, and by a top/front light (25 W warm light), while the rest of the enclosure was dimly illuminated. In the running wheel, chicks could easily invert the direction of movement.

### Test procedure

Chicks were individually placed in the running wheel facing the long side of the enclosure, so that they could see both stimuli with their lateral eyes, and tested for their spontaneous preference to walk toward the stuffed hen and scrambled hen. Chicks could operate the wheel by walking towards each stimulus, while the distance run (in metres) was recorded by an automated system connected to the wheel. The distance run was checked every 5 minutes (minute 5, 10, 15, 20, 25, 30), for 30 minutes overall.

### Statistical analysis

To assess spontaneous preferences for the stuffed hen independently from motor activity, for each chick we calculated a relative **hen preference** index, adjusted for its overall distance run as:

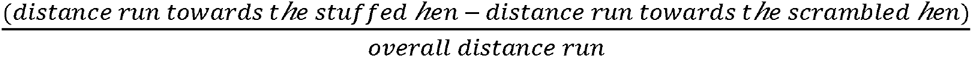

where 0 indicates no preference, 1 a complete preference for the stuffed hen and -1 a complete preference for the scrambled hen.

Because the data did not fit linear or commonly used non-linear models, we pursued non-parametric statistics. Significant deviations between breeds (Padovana, Polverara, Robusta) were assessed using the Kruskall Wallis test for the overall session (30 minutes). To establish the presence of significant deviations from the chance level (0), that could indicate a significant preference for the tested stimuli, we used one sample Wilcoxon Signed rank tests. Post-hoc comparisons between breeds and time points were conducted using Mann Whitney U test and Wilcoxon Signed rank test.

We assessed differences in the overall **motor activity** between breeds comparing the overall distance run (in metres) irrespectively of the stimulus chosen, for the overall session (30 minutes) and for the six time periods. Due to its data distribution, this variable was analysed using non-parametric statistics. To assess whether the strength of the preferences for the stuffed hen depended on the amount of motor activity, we calculated a Spearman’s rank correlation between the preference and motor activity. Exploratory and statistical analyses were performed with the R software (version 3.1.2).

## Results and discussion

### Hen preference

To investigate the predisposition for approaching the stuffed hen *vs.* the scrambled hen, we used the relative hen preference index, which indicates the relative preference for the stuffed hen *vs.* the scrambled hen independently from the amount of activity in the wheel (see Figure 3). In fact motor activity could be affected by differences between breeds other than their predisposed preferences, such as motor development. A Kruskal Wallis test with Dunn’s multiple comparisons (Bonferroni correction) showed a significant effect of Breed (chi-squared_2_=7.167, p=0.028), with significant differences between Padovana and Polverara (Z=−2.219, p=0.040), Polverara and Robusta (Z=−2.390, p=0.025) and no significant difference between Padovana and Robusta (Z=−0.189, p=1), see Figure 3A.

**Figure 3.**
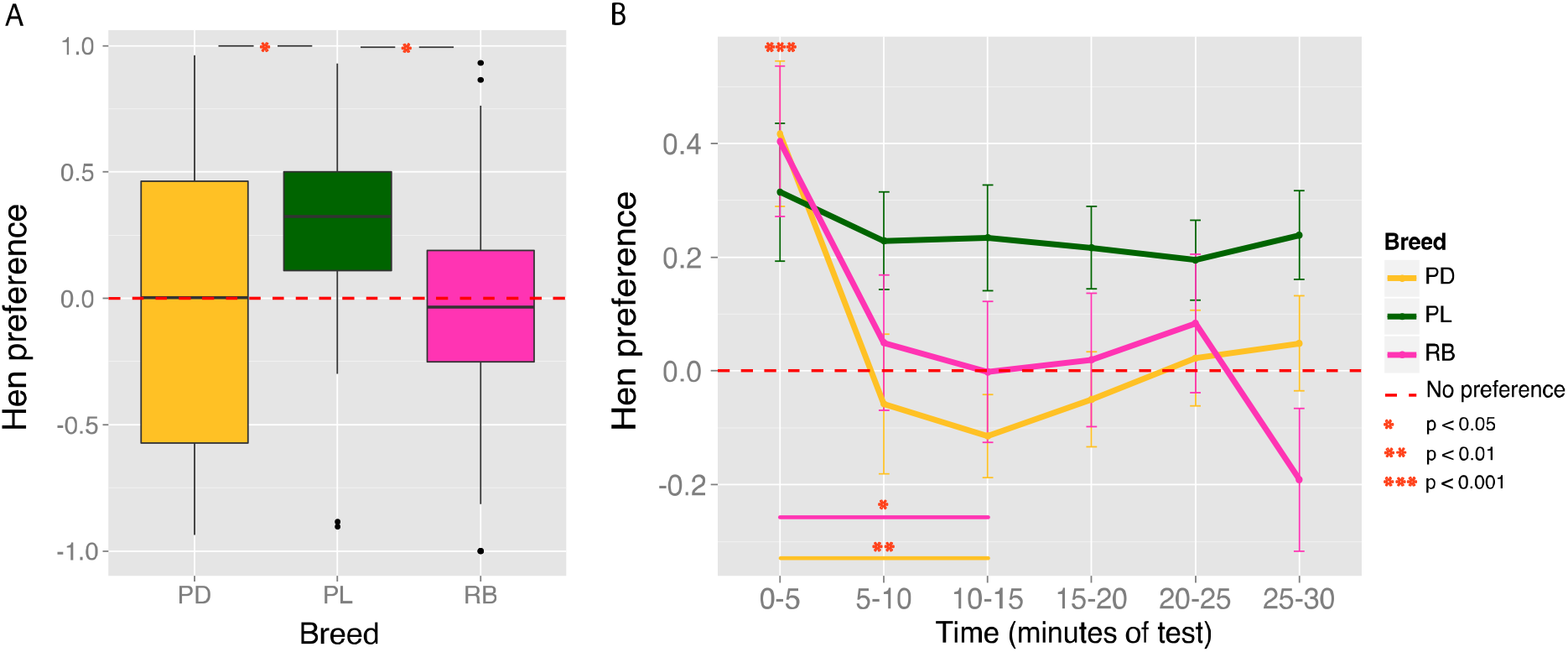
(A) Hen preference across the entire 30 minutes of test by Breed, as: (distance to stuffed hen - distance to scrambled hen)/overall distance run. Boxplots show median and quartiles. (B) Hen preference by Breed in Time (every 5 minutes of test): means +/- standard error of the mean are plotted.

Considered *per se*, this result would suggest not only differences between breeds in the predisposed preference for the stuffed hen, but also a lack of predisposition in the Padovana and Robusta breeds (Figure 3A?). Nevertheless, when analysing the performance of the three breeds across time, as shown in Figure 3B, the scenario appears much different: at the very beginning of their visual experience (minutes of test 0-5), there was no significant difference between breeds (chi-squared_2_=2.856, p=0.240) but we observed an overall significant preference for the stuffed hen (V=3120, p<0.001). Hence, in their first moments of life all breeds were attracted by the stimulus that presented more animacy cues, showing a predisposition for the stuffed hen over the scrambled hen. Differences between breeds emerged in the continuation of the experiment, and maximized after 10-15 minutes of exposure, with a significant reduction of preference for the hen in the Padovana and Robusta breeds (V=330, p=0.004 and V=205, p=0.042, respectively), while the Polverara breed maintained the same preference (V=245.5, p=0.552).

How can fluctuations in preference for the stuffed hen be explained from an ethological point of view? At least two mechanisms can be responsible for it. First, in the wild, chicks can usually approach the naturalistic stimuli to which they direct their affiliative responses, and receive visual, tactile and acoustic feedback [28,29]. This feedback is very important to maintain proximity with the stimulus and induce the filial imprinting process. Filial imprinting is a fast learning process that enables chicks to learn the features of their social partners and to restrict their affiliative responses to them by mere exposure [reviewed in 30,31,32]. Not only movement and auditory signals [28,33–35] of the object increase its attractiveness and effectiveness as imprinting object, but the interaction with the mother induces greater preferences for it, compared to experience with a moving stuffed model [28]. Hence, a first explanation for the decrease of the predisposed preference for the stuffed hen is the absence feedback from the stimulus. Second, chicks search exposure to novel stimuli before the filial imprinting process is terminated, likely to form a more comprehensive representation of it that enables recognition from novel points of view [36–38]. Consistent evidence has shown that, especially in the early stages of imprinting, the tendency to approach the familiar object can be temporarily reversed [e.g. 31,37,39,40], and that chicks actively search for novel aspects of the imprinting object [41]. Given that our test is performed at the very beginning of the imprinting process, a change in preferences during the test after a first orienting response towards the predisposed stimulus is consistent with the ethological needs of the filial imprinting process [31,37].

### Motor activity

To check whether the tested breeds differ in early motor activity, and if a connection between motor activity and predisposed preferences exists, we measured the distance run in the wheel and explored the correlation between motor activity and the relative hen preference index discussed above. A Kruskal Wallis test with Dunn’s multiple comparisons (Bonferroni correction) showed a significant effect of Breed (chi-squared_2_=31.563, p<0.001), with significant differences between Padovana and Robusta (Z=−24.494, p<0.001), Polverara and Robusta (Z=−5.198, p<0.001) and no significant difference between Padovana and Polverara (Z=0.674, p=0.75), see Figure 4A.

**Figure 4.**
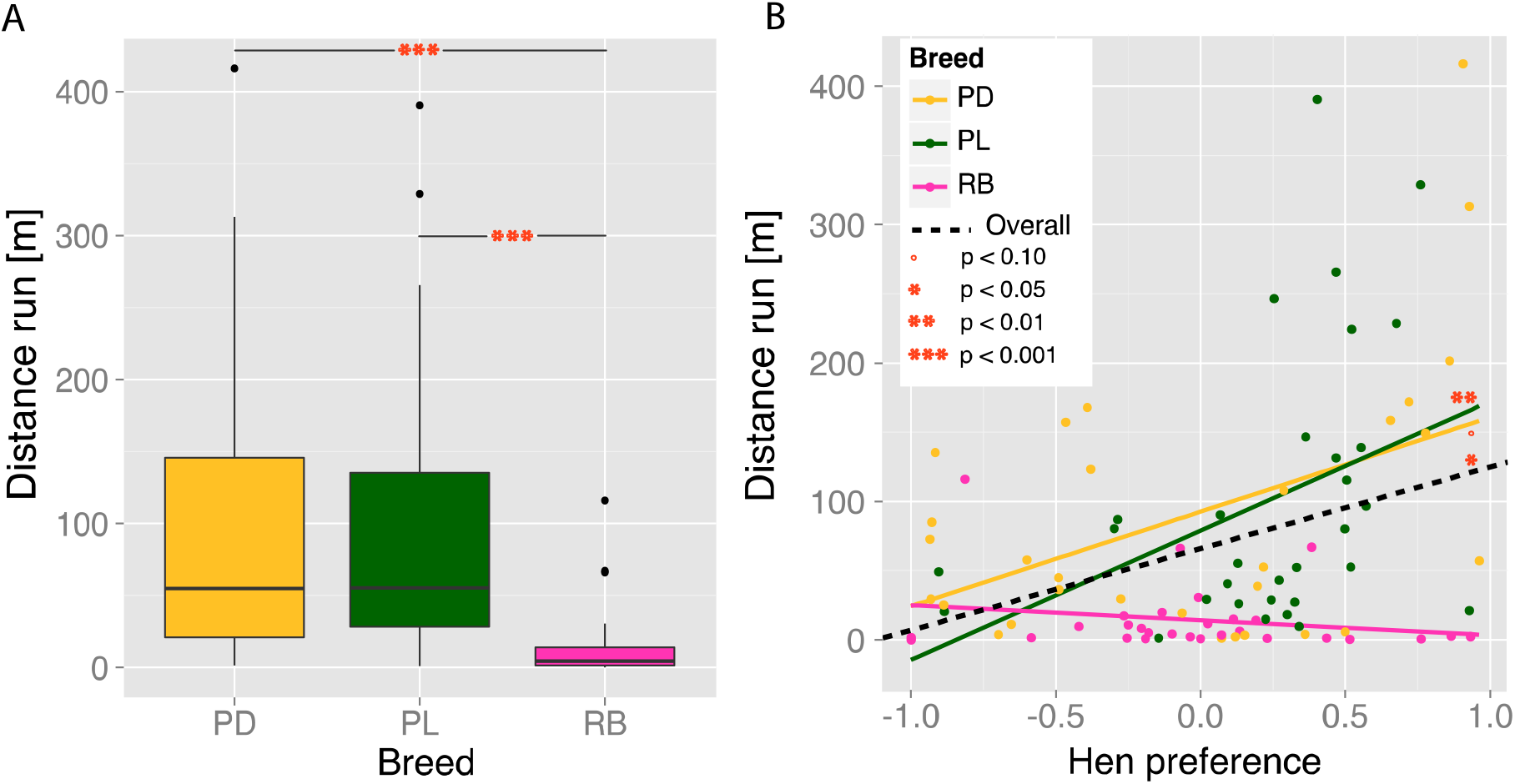
(A) Motor activity (distance run in metres) across the entire 30 minutes of test by Breed. Boxplots show median and quartiles. (B) Relation between Motor activity and Hen preference by Breed.

Interestingly, the differences in motor activity between breeds could dissociate from differences in hen preference. In particular, Padovana and Robusta breeds did not differ in their stuffed hen preference but greatly differed in motor activity. This strongly suggests that the differences in the predisposed preference for the stuffed hen do not simply reflect the motor activity or motor development of the different breeds.

Nevertheless, it would still be possible that motor activity is a proxy for affiliative motivation. To check for this hypothesis, we computed the correlation between the overall motor activity and the hen preference index: considering all breeds, we found a significant positive correlation (Spearman’s ρ=0.254, p=0.016), which shows that, overall, chicks with a stronger stuffed hen preference also exhibited a higher motor activity (Figure 4B). When considering single breeds, this correlation was significant for the Polverara breed (ρ=0.461, p=0.010), not significant – but close to significance – for the Padovana breed (ρ=0.315, p=0.090), and not significant for the Robusta breed (ρ=−0.11, p=0.538). In the first period of the test (minutes 0-5), before any imprinting took place, we observed an overall significant positive correlation between hen preference and motor activity (ρ=0.318, p=0.002), which was close to significance for the Polverara breed (ρ=0.330, p=0.070), significant for the Padovana breed (ρ=0.517, p=0.003) and not significant for the Robusta breed (ρ=0.073, p=0.706). Hence, although motor activity *per se* is not the trigger of predisposed preferences – otherwise we would have observed in Robusta chicks the same association between motor activity and stuffed hen preference observed in other breeds –, motor activity is associated with predisposed preferences. This finding suggests that predisposed preferences enhance affiliative responses.

### General discussion

Human neonates and chicks of the domestic fowl share biases to prefer face-like stimuli [2,12] and other cues associated with animate objects [reviewed in 1,13,14], such as biological motion [5,6,42], changes of speed [8] and self-propulsion [9,10]. Individual variability in these predispositions has been observed in both species [2,12,18,43], and in human neonates it is linked to high risk of developing Autism Spectrum Disorders [43]. Understanding whether individual variability in early predispositions has a genetic component would be of primary interest for biomedical research. The spontaneous preferences of chicks for a stuffed hen *vs.* a stimulus in which the head configuration had been disrupted have been systematically reported [12,17,18,27]. We used genetically different chicken breeds [16], which have been maintained genetically isolated for at least eighteen years [22], to identify the presence of segregating variability in the predispositions of chicks in approaching a stuffed hen. Overall, in visually naïve chicks of all three tested breeds (Padovana, Polverara and Robusta maculata) we observed the same initial preference for the predisposed stimulus, suggesting that the direction of the initial preference might be genetically fixed across the tested breeds or at the species level, given that the same direction of preference had been previously observed in broilers of different strains [12,13,17,18]. Few minutes after the first exposure though, striking differences emerged between breeds, that could indicate early different strategies of dealing with affiliative objects: while the Polverara breed maintained a constant preference for the entire test, the Padovana and Robusta breeds progressively explored the alternative stimulus more. This second strategy, in line with the motivation of chicks to be exposed to novel stimuli at the beginning of the filial imprinting process [37,39–41], might help chicks in looking for responsive parental objects and in forming a more structured representation of the mother hen. When analysing the connection between predisposed preferences and motor activity we identified a partial dissociation: the initial preference did not depend on motor activity (the preference for the stuffed hen was present in highly and less mobile chicks), but overall a positive correlation between motor activity and hen preferences was present. This result corroborates the hypothesis that the function of predispositions for animacy cues is to orient the individual towards the social partners, and, in the case of domestic chicks, this can be the basis for the strong attachment mechanism of filial imprinting, which implies approaching responses [28,30,33]. Given that all tested breeds have been farmed in the same way for decades, and that all eggs and chicks have been exposed to the same treatments, the observed behavioural differences indicate the presence of inherited variability in early social predispositions. This study paves the way to further genomic investigation of the variability in predisposed preferences for animate objects in the chick as a model system. Understanding the genetic basis of predispositions for animacy cues and its individual variability, it might have a crucial importance for translational studies on developmental pathologies, such as Autism Spectrum Disorders [43]. The use of chicks as system model is particularly suitable not only for the ease of handling and controlling precocial special species until the moment of test and for the established parallels between human newborns and chicks [1], but also for the mounting evidence on the neurobiological basis of spontaneous predispositions [7,18,27] and the availability of genomic tools [44–47] and controlled populations with segregating variation [16,46].

## Authors’ contributions

EV and GV conceived the experiments; EV, ADZ and GV designed the experiments; IF and EV conducted the experiments; GB maintained the chicken breeds; EV analysed the data; EV drafted the manuscript; EV, IF, GB, ADZ and GV wrote the manuscript. All the authors gave final approval for publication.

## Acknowledgements

We thank Tommaso Pecchia for help with the experimental apparatus.

